# Bacteria sense virus-induced genome degradation via methylated mononucleotides

**DOI:** 10.1101/2025.11.05.686725

**Authors:** Ilya Osterman, Bohdana Hurieva, Sarit Moses, Alla H. Falkovich, Maxim Itkin, Sergey Malitsky, Erez Yirmiya, Rotem Sorek

## Abstract

Phages often degrade the genome of their bacterial host to individual nucleotides and use these nucleotides to build their own genome. In this study, we describe a bacterial defense system that directly senses phage-mediated host genome degradation. This system, called Metis, aborts phage infection once it detects the accumulation of the modified mono-nucleotide N⁶-methyl-deoxyadenosine monophosphate (m⁶dAMP). As methylation of deoxy adenosines occurs only in the context of the DNA polymer, intracellular accumulation of m⁶dAMP serves as a definitive signal that the host genome has been degraded to its individual constituents. In type I Metis, sensing of m⁶dAMP activates an NAD^+^ diphosphatase, leading to rapid NAD⁺ depletion and cessation of the infection process; while the effector in type II Metis is a transmembrane-spanning protein whose toxicity is triggered in response to the modified mono-nucleotide. We further show that Metis defense depends on endogenous DNA methylases, and that phages can escape Metis via mutations that inactivate phage-mediated host genome degradation. Our results demonstrate how molecular byproducts released during virus-induced cell exploitation can be used as specific danger signals that trigger host immunity.

## Introduction

Viruses, being obligatory parasites, exploit host cell resources to replicate. When lytic phages infect bacterial cells, one of the most common forms of resource exploitation is the complete degradation of the host DNA and utilization of the emerging deoxynucleotides as building blocks to construct the phage genome. *Escherichia coli* phage T4, for example, degrades host DNA into individual deoxynucleotides already starting 3-5 minutes after the onset of infection at 37°C^1,2^, and coliphage T7 initiates host genome degradation eight minutes after initial infection^3^. Phage T5 also completely degrades host DNA early upon infection^4^, although this phage synthesizes its own nucleotides and does not rely on the degraded host DNA to build its genome^5^. Here, we report the discovery of a bacterial defense system that senses methylated nucleotides released as byproducts of host genome degradation.

DNA methylation is a common epigenetic modification in organisms across the tree of life^6^. In bacteria, one of the most abundant DNA modifications is methylation of adenine in the N6 position of the base. The model organism *E. coli* encodes a DNA adenine methyltransferase (Dam) that methylates adenines in the context of 5′-GATC-3′ double-stranded DNA sequence motifs^7^. Methylated GATC motifs regulate replication initiation and DNA mismatch repair in *E. coli* and other Proteobacteria^8,9^. Adenine methylation is also common as part of restriction-modification systems in many bacteria, where the restriction sequence motif is methylated in the host genome to prevent cleavage of self DNA^10^. DNA adenine methylation always occurs in the context of the DNA polymer, such that adenines are methylated only when they are part of the polymerized DNA chain^11^.

In the current study we describe Metis, a defense system named after the Titan-goddess of wisdom and deep thought in Greek mythology. This system specifically recognizes N⁶-methyl-deoxyadenosine monophosphate (m⁶dAMP), a mononucleotide that accumulates in the cell when phage nucleases degrade methylated host DNA. We show that m⁶dAMP directly activates a Metis-encoded NAD^+^ diphosphatase effector, leading to NAD⁺ depletion via an enzymatic activity not previously known to participate in anti-phage immunity, which arrests the infection process. We also discover type II Metis systems, in which m⁶dAMP activates a membrane-spanning effector that likely impairs membrane integrity. We find that Metis provides protection against multiple unrelated lytic phages, including T2, T4, T5, T6 and T7, and that Metis defense can be evaded when phage genes responsible for host DNA degradation are inactivated. Although the infected bacterium is essentially dead once its genome has been degraded by the phage, the Metis system prevents replication of phages in the infected cell, thus protecting the nearby cells from spread of the phage epidemic.

## Results

### A two-gene defense system protects bacteria from lytic phages

We studied the anti-phage activity of a defense system encoding two proteins with predicted metallophosphatase activities (Figure 1A). The first protein, called here MisA, has a Pfam annotation of calcineurin-like phosphoesterase (Pfam accession PF00149), while the second protein, MisB, is annotated as a haloacid dehalogenase-like phosphatase (Pfam accession PF13419). An operon with a similar gene composition was recently implicated with an anti-phage function in a machine-learning-based screen aimed at discovering new defense systems^12^. We synthesized and cloned three such systems from *E. coli* strains 401675, 402837, and E308, and found that all three systems conferred protection against the lytic phages T2, T4, T5, T6, and T7 (Figure 1B, S1A). As the system from *E. coli* 401675 conferred the strongest defense, we used this system for further experiments *in vivo*. Infection experiments in liquid culture showed that the system protects the culture from phage-mediated collapse at low multiplicity of infection (MOI), while at high MOI the culture collapsed even when expressing the defense system (Figures 1C, S1B, S1C). These results suggest that cells encoding the two-gene operon do not survive infection, but prevent the phage from producing viable progeny. This defense system is henceforth denoted Metis.

**Figure 1.**
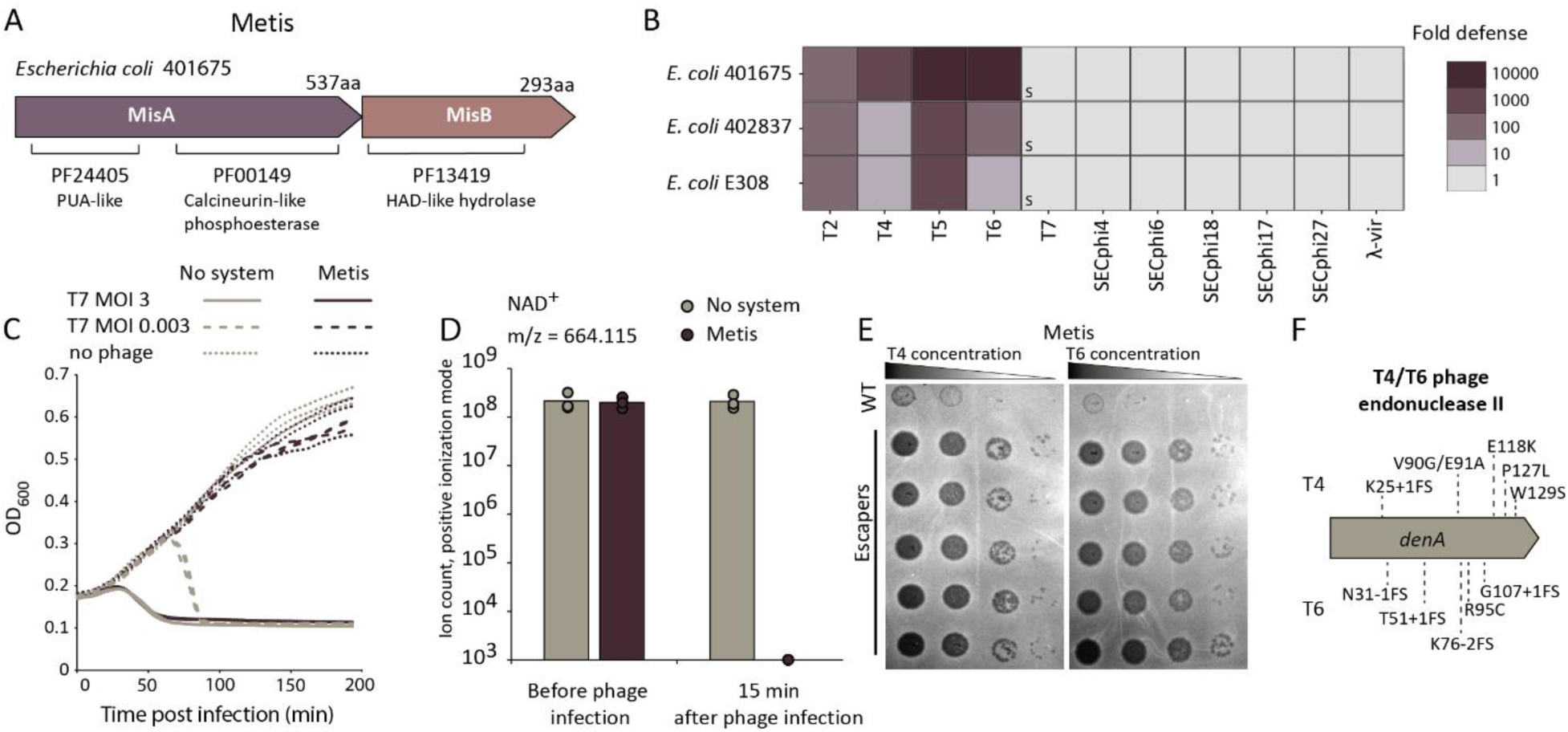
Metis is an anti-phage defense system that degrades NAD⁺. A. Schematic representation of protein domain composition in the Metis system. B. Metis systems protect against phages. Systems from three *E. coli* strains were expressed in *E. coli* MG1655, a strain that naturally lacks Metis. Fold defense was quantified by serial dilution plaque assays, comparing the efficiency of plating of phages on the system-containing strain to the efficiency of plating on a control strain that lacks the systems and contains an empty vector instead. Data represent an average of three replicates. PFU quantification for individual replicates is presented in Figure S1A. ‘‘s’’ designates a marked reduction in plaque size. C. Growth curves of *E. coli* MG1655 cells expressing the Metis system from *E. coli* 401675 or an empty vector, infected by phage T7 at an MOI of 0.003 or 3 (or 0 for uninfected cells). Data from three biological replicates are presented as individual curves. D. NAD⁺ is depleted in cells expressing Metis during infection with T7 phage. Cells were infected at MOI = 3, measurements taken 15 min post infection. Presented are untargeted LC-MS-derived ion count data; bars represent the mean area under the curve of three experiments, with individual data points overlaid. E. Plaque assays showing T4 and T6 mutants that escape Metis defense. Data are representative of three biological replicates of phages infecting *E. coli* MG1655 cells that express the Metis system from *E. coli* 401675. PFU quantification is presented in Figures S1D and S1E. F. The *denA* gene is mutated in T4 and T6 phages that escape Metis defense. Detected mutations are annotated; FS, frameshift.

A large diversity of bacterial defense systems is known to inhibit phage replication by manipulating the intracellular pool of metabolites essential for core cellular processes^13^. To examine whether the Metis system affects the concentration of essential cellular metabolites, we extracted cell lysates 15 minutes after infection by phage T7 and performed liquid chromatography and mass spectrometry (LC-MS) analyses. We found that in cells expressing the Metis defense system, the molecule nicotinamide adenine dinucleotide (NAD⁺) was completely depleted during infection (Figure 1D). NAD^+^ depletion was not observed in cells lacking Metis, suggesting that NAD^+^ elimination is the consequence of the system’s activity (Figure 1D). It was previously shown that NAD^+^ depletion is an efficient modality of anti-phage defense, and that many bacterial defense systems, including CBASS^14^, Pycsar^15^, Thoeris^16^, and prokaryotic argonautes^17^, deplete NAD^+^ to inhibit phage propagation. Notably, infected cells expressing the Metis defense system also exhibited partial or full depletion of several dNTPs and NTPs in addition to NAD^+^ (Figure S2).

We were able to readily isolate T4 and T6 phage mutants that escaped Metis-mediated defense (Figure 1E, S1D, S1E). Sequencing the genomes of five such escaper T4 phages and five T6 escapers revealed that all these phages were mutated in *denA*, a phage gene that encodes the endonuclease II enzyme responsible for the initial step of host genome degradation^18^ (Fig 1F). Many of these mutations involved frame shifts, suggesting that the mutated gene does not produce a viable protein product. It was previously shown that T4 phages deficient in endonuclease II do not degrade host DNA, but can produce viable progeny when grown in laboratory conditions^19^. As all phages sensitive to Metis defense degrade the host genome as part of their life cycle, these data imply that the Metis system might somehow monitor the integrity of the host DNA and becomes active when host DNA is degraded by phage.

### m⁶dAMP activates NAD⁺ hydrolysis by MisA

Given our hypothesis that Metis defense is activated in response to bacterial genome degradation, we set out to seek the exact molecular signal that triggers the system. The N-terminal domain of MisA provided an initial clue that the signal might involve nucleotides with a modified base. This domain is structurally similar to the PUA domain^20^, which is known to be involved in the recognition of modified DNA and RNA bases (Figure 1A)^21^. We therefore hypothesized that modified deoxynucleotides released from the host DNA during DNA degradation might form the Metis-activating signals.

In *E. coli* MG1655, genomic DNA is primarily modified by the Dam enzyme, which installs methyl residues on adenines in GATC sites^22^, and the Dcm methylase, which methylates cytosine residues in the context of CCAGG and CCTGG sequences^23^. The Metis system was active when expressed in *Δdcm E. coli* cells, but defense was completely abolished when Metis was expressed in *Δdam* strains, suggesting that host Dam activity is essential for Metis function (Figure 2A).

**Figure 2.**
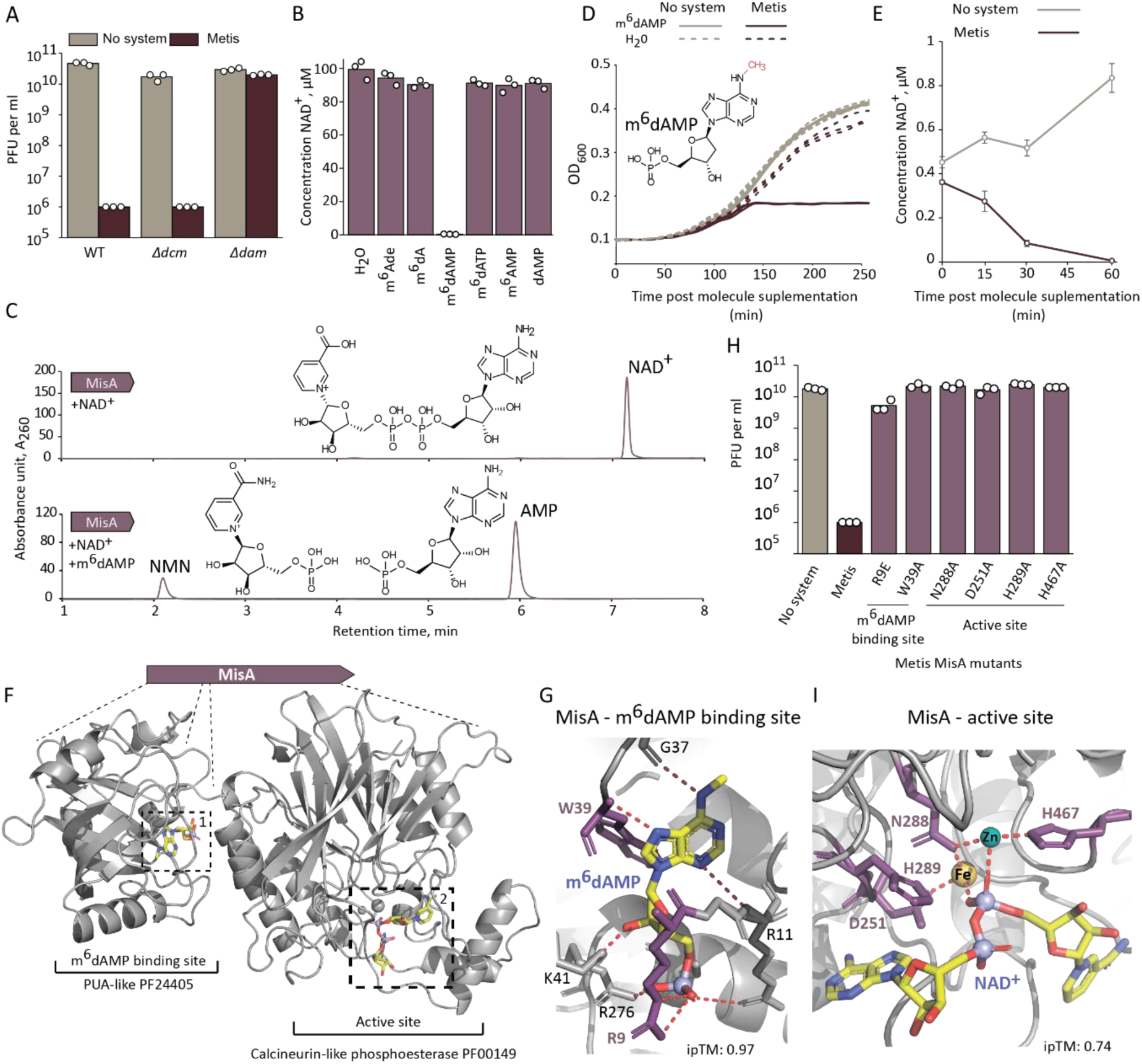
MisA is activated by m⁶dAMP to cleave NAD^+^. A. Metis is not active in cells in which *dam* is deleted. Data represent PFUs per milliliter of T5 phage infecting control cells (no system) and cells expressing Metis, transformed to WT *E. coli*, Δ*dam* strain or Δ*dcm* strain. Bar graphs represent the average of three independent replicates, with individual data points overlaid. B. MisA degrades NAD^+^ *in vitro* in the presence of m^6^dAMP. 100µM NAD^+^ was incubated with 1 µM purified MisA and 10 µM synthetic adenine variants for 120 minutes, and NAD^+^ levels were measured by the NAD/NADH-Glo biochemical assay. Average of three replicates with individual data point overlaid. C. HPLC analysis of the products of NAD^+^ degradation by MisA in the presence or absence of 10 µM m⁶dAMP. Products of cleavage were identified by comparing the peaks with the retention time of chemical standard molecules (Figure S3B). D. m⁶dAMP is toxic to Metis-expressing cells. Shown are growth curves of *E. coli* MG1655 cells expressing the Metis system from *E. coli* 401675 or an empty vector after addition of 50 µM m⁶dAMP to the growth media. Three replicates are presented as individual curves. The chemical composition of m^6^dAMP is presented above the curves. E. NAD^+^ concentration in lysates obtained from cells expressing Metis, or control cells with an empty vector instead (no system). Samples were analyzed 0,15,30 and 60 minutes after adding 50 µM m⁶dAMP to the growth media. Average of 3 replicates, error bars represent standard deviation. F. AlphaFold3-predicted structure of MisA from *E. coli* 401675 in complex with m⁶dAMP, NAD^+^, Zn^2+^ and Fe^3+^. Binding sites of m⁶dAMP (1) and NAD^+^ (2) are shown in dashed boxes. G. Close-up view of the predicted m⁶dAMP binding site, ipTM of the complex of MisA with m⁶dAMP is presented. Red dashes indicate predicted hydrogen bonding interactions. H. Mutations in MisA abolish defense against phage. Data represent PFUs per milliliter of T5 phage infecting cells that express WT Metis or Metis mutated in *misA* in the indicated residues. Bar graphs are the average of three independent replicates, with individual data points overlaid. I. Close-up view of the predicted NAD^+^ binding site. ipTM of MisA co-folded with m⁶dAMP, NAD^+^, Zn^2+^ and Fe^3+^ is presented. Amino acids mutated in this study are presented in purple. Red dashes indicate predicted hydrogen bonding interactions.

As Dam methylates adenines on the N^6^ position on the DNA polymer, we suspected that the Metis-activating signal would involve methylated adenines. To test this hypothesis, we purified MisA for *in vitro* biochemical experiments. As MisA from *E. coli* 401675 did not purify well, we instead used a homolog from *E. coli* 402837 Metis, which also defends against phages (Figure 1B). Purified MisA was incubated with a series of modified and non-modified adenine variants, and its ability to hydrolyze NAD^+^ was evaluated. MisA exhibited strong NAD⁺-degrading activity *in vitro* in the presence of the monophosphorylated nucleotide m⁶dAMP (Figure 2B). No NAD^+^ degradation was observed when MisA was incubated with related molecules, including non-methylated dAMP, the methylated ribonucleotide m⁶AMP, the non-phosphorylated nucleoside m⁶-deoxyadenosine, or the methylated base m⁶-adenine, demonstrating that m⁶dAMP is the specific signal that activates MisA (Figure 2B).

HPLC analysis revealed that MisA hydrolyzes NAD⁺ *in vitro* to produce nicotinamide mononucleotide (NMN) and AMP, showing that MisA is an NAD^+^ diphosphatase that cleaves the bond between the two phosphates of the NAD^+^ molecule (Figure 2C). These results were consistent with LC-MS data from in vivo samples, in which NMN levels increased by two orders of magnitude in infected, Metis-expressing cells (Figure S2). Incubation of MisA with m⁶dAMP and ATP did not cause ATP degradation, suggesting that NAD^+^ is the primary target of MisA, and that the reduced levels of ATP detected *in vivo* may be a secondary effect of MisA-mediated NAD^+^ depletion (Figure S3A).

To test whether m⁶dAMP can activate Metis toxicity *in vivo*, we added synthetic m⁶dAMP to growth media and monitored bacterial proliferation. Supplementation of 50 µM m⁶dAMP to the media was sufficient to cause growth arrest in cells carrying Metis, but not in control cells lacking the system (Figure 2D). NAD⁺ measurements in lysates extracted from cells following incubation with m⁶dAMP confirmed depletion of NAD⁺ in Metis-containing cells, even in the absence of phage infection (Figure 2E). These results demonstrate that m⁶dAMP activates Metis to deplete NAD^+^ both *in vitro* and *in vivo*.

To gain further insight into the molecular basis for m⁶dAMP recognition by MisA, we used AlphaFold3^24^ (AF3) to co-fold MisA and m⁶dAMP. AF3 modelled m⁶dAMP within the N-terminal PUA-like binding pocket with very high confidence (ipTM = 0.97) (Figures 2F, 2G). Point mutations in residues predicted to be in contact with m⁶dAMP in the binding pocket abolished defense, supporting the prediction that these residues participate in m⁶dAMP recognition (Figure 2H). As expected, the AF3 model placed NAD⁺ within the C-terminal metallophosphatase catalytic site of MisA, and predicted that NAD⁺ catalysis is coordinated by Zn²⁺ and Fe²⁺ ions, as known for metallophosphatases of this family^25^ (Figure 2I). Mutations in predicted catalytic site residues abolished Metis defense (Figure 2H).

### MisB prevents system toxicity by degrading basal levels of m⁶dAMP

Although MisA is frequently encoded as part of a *misA*-*misB* operon (Figure 1A), we were surprised to find that cells expressing MisA alone were resistant to phage infection, suggesting that MisA encompasses the defensive capacity of the system and that MisB is not essential for defense (Figure 3A). Nevertheless, the strain expressing MisA alone grew slower than the strain encoding the full system, suggesting that MisB is necessary to prevent MisA toxicity in the absence of phage (Figure 3B). Indeed, basal NAD⁺ levels in cells encoding MisA alone were lower than those measured in WT cells or in cells expressing both MisA and MisB, explaining the slower growth and showing that MisA is residually active in the absence of MisB (Figure 3C).

**Figure 3.**
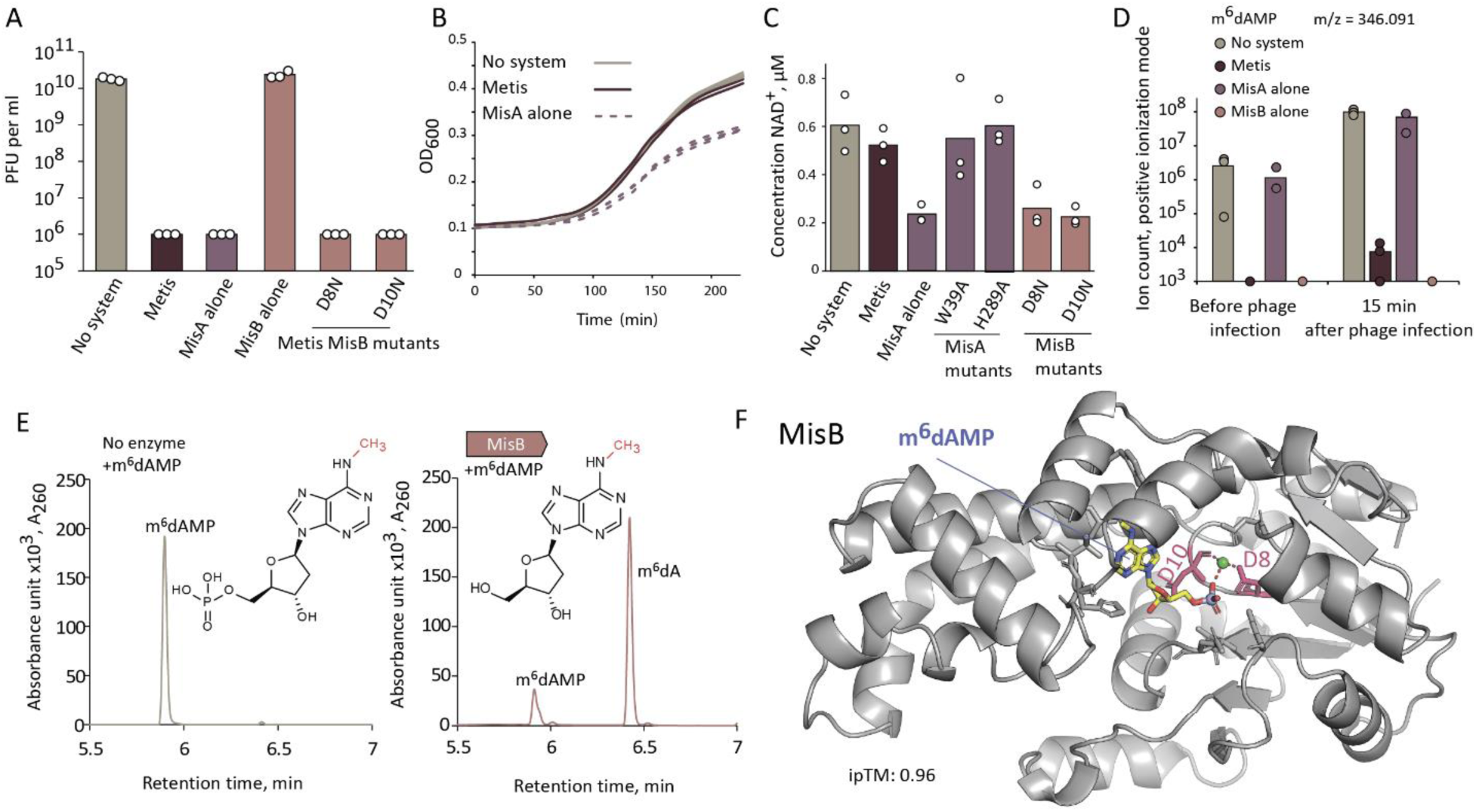
**MisB degrades residual m⁶dAMP and prevents Metis toxicity**. A. MisA alone or Metis with mutated MisB, but not MisB alone, protects bacteria from phage infection. Data represent PFUs per milliliter of T5 phage infecting cells that express the indicated system, measured via serial dilution plaque assays. Bar graphs represent the average of three independent replicates, with individual data points overlaid. B. In the absence of MisB, MisA causes growth retardation. Growth curves of *E. coli* MG1655 cells expressing Metis, MisA alone, or an empty vector (no system). Data from three replicates are presented as individual curves. C. NAD^+^ concentration measured in lysates extracted from cells expressing WT Metis, MisA alone, or the Metis system with point mutations in MisA or MisB. Cells were collected from the experiment shown in panel B after 100 minutes of growth. Bar graphs represent the average of three independent replicates, with individual data points overlaid. D. MisB eliminates basal levels of m⁶dAMP. Cells were collected before infection and 15 minutes following infection with phage T7 (MOI = 3). Presented are LC-MS ion count data; bars represent the average area under the curve in three experiments, with individual data points overlaid. E. LC-MS analysis of m⁶dAMP degradation by MisB in the presence of Mg^2+^ ion. The product was identified by comparing the observed peak with the retention time and m/z signals of m^6^dA chemical standard (Figure S4C). F. AlphaFold3-predicted structure of MisB from *E. coli* 401675 in complex with m⁶dAMP and Mg^2+^ (ipTM = 0.96). Mutated residues are highlighted in pink; Mg²⁺ is shown in green. Red dashes indicate predicted hydrogen bonding interactions.

As MisA activity is triggered by m⁶dAMP, we hypothesized that low levels of m⁶dAMP are present in the cell even in the absence of full genome degradation. Some m⁶dAMP can potentially be generated through host exonuclease activity during DNA mismatch repair or during RecBCD-mediated DNA processing^26^. LC-MS analysis confirmed the presence of basal levels of m⁶dAMP in lysates from Metis-lacking cells prior to phage infection (Figure 3D). However, no m⁶dAMP was detected in lysates from non-infected cells expressing MisB, either alone or in the context of Metis (Figure 3D). These results suggest that the role of MisB is to degrade basal levels of m⁶dAMP to prevent activation of MisA before phage infection.

Given that MisB is annotated as a Mg²⁺-dependent phosphatase from the HAD superfamily, we hypothesized that MisB inactivates m⁶dAMP by removing the phosphate from the nucleotide. Incubation of purified MisB from *E. coli* 401675 with m⁶dAMP confirmed this hypothesis, and showed that MisB efficiently dephosphorylates the molecule to generate m^6^dA (Figure 3E), a molecule that cannot activate MisA (Figure 2B). Co-folding of MisB with m⁶dAMP using AF3 yielded a high confidence model in which m⁶dAMP is placed in the active site pocket of the MisB phosphatase (Figures 3F), which contains two aspartic acid residues typical to HAD phosphatases^27^. Mutations in these residues led to reduced growth rate and reduced NAD⁺ levels in the absence of phage, confirming that the catalytic activity of MisB is necessary for toxicity prevention (Figures 3C, S4A, S4B).

### Type II Metis encodes a transmembrane effector that senses m^6^dAMP

Homologs of *misB* in diverse bacteria are frequently encoded next to *misA* homologs, confirming that these two genes function together (Figure 4A, Table S1). However, our analyses show that in many bacterial genomes *misB* homologs are not in the vicinity of a *misA* gene, but instead appear in an operon with a gene lacking an annotated protein domain. This gene, which we denote *misC*, encodes a protein with three predicted membrane-spanning helices (Figure 4A, 4B). A MisB-MisC operon cloned from *Phyllobacterium sp.* UNC302MFCol5.2 and expressed in *E. coli* MG1655 conferred anti-phage defense (Figure 4C). Notably, an operon from *E. coli* strain 2862600, encoding homologs of MisC and MisB, was previously shown to defend against phage T7^28^.

**Figure 4.**
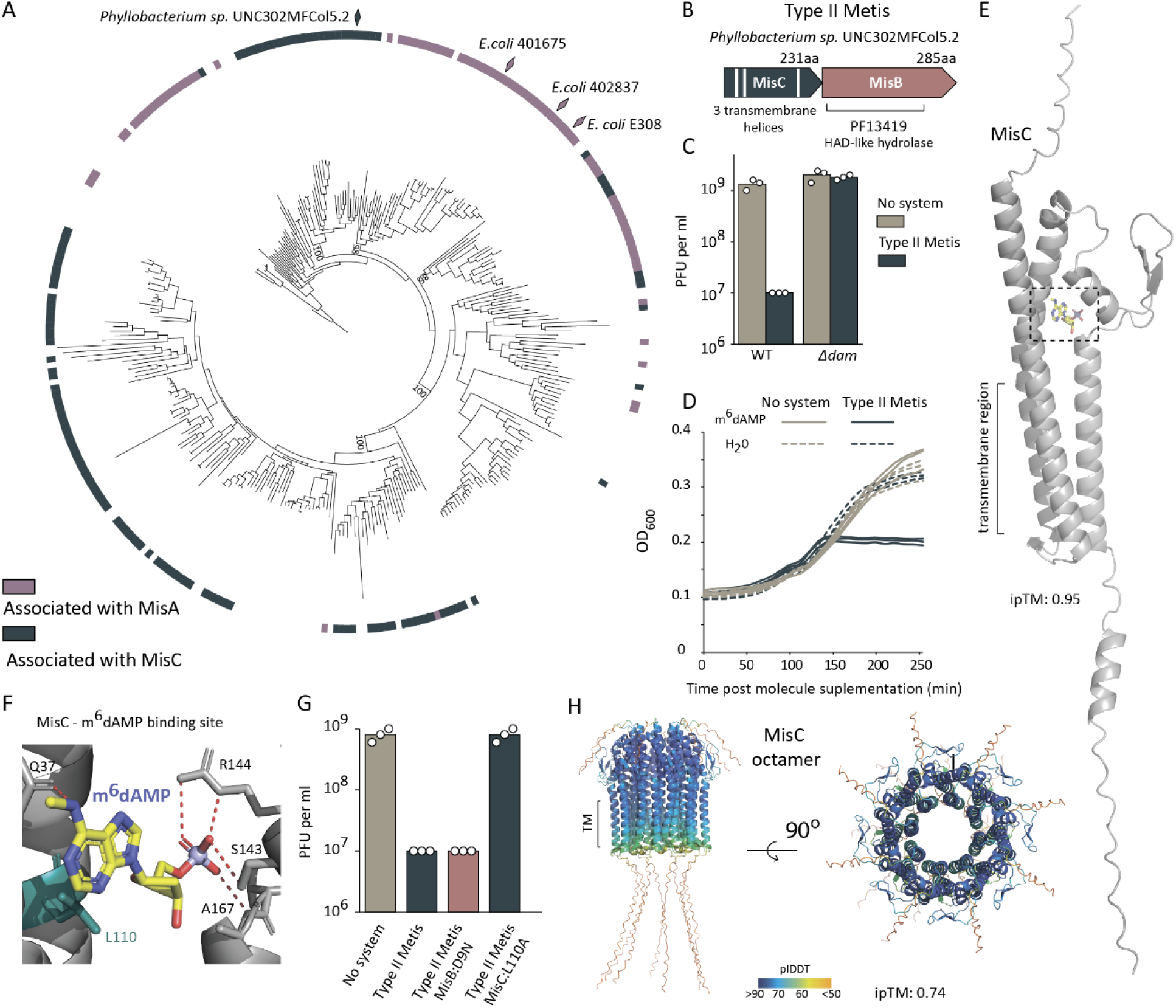
Type II Metis. A. Phylogenetic analysis of the MisB protein family in bacterial genomes. The outer ring indicates whether MisB is associated with MisA or MisC. Bacteria carrying systems experimentally validated in this work are marked on the tree. Bootstrap values are shown for major clades. B. Schematic representation of the type II Metis system. C. Type II Metis is not active in cells in which *dam* is deleted. Data represent PFUs per milliliter of T6 phage infecting control cells (no system) and cells expressing type II Metis from *Phyllobacterium sp.* UNC302MFCol5.2, transformed to WT or Δ*dam E. coli* strains. Bar graphs represent the average of three independent replicates, with individual data points overlaid. D. Growth curves of *E. coli* MG1655 cells expressing type II Metis or cells with an empty vector, after addition of 500 µM m⁶dAMP to the growth media. Data from three replicates are presented as individual curves. E. AlphaFold3-predicted structure of MisC from *Phyllobacterium sp.* UNC302MFCol5.2 in complex with m⁶dAMP (ipTM = 0.95). Predicted binding site of m⁶dAMP is shown in a dashed box. F. Close-up view of the predicted m⁶dAMP binding site in MisC. Amino acid selected for mutagenesis is presented in dark green. G. Point mutations in MisC, but not in MisB, abort type II Metis defense. Data represent PFUs per milliliter of T6 phage infecting cells that express WT Metis or Metis genes mutated in the indicated residues. Bar graphs are the average of three independent replicates, with individual data points overlaid. H. AlphaFold3-predicted structure of MisC octamer, ipTM=0.74. Colors represent pLDDT scores.

To test whether the *Phyllobacterium* MisC-MisB operon shares functional features with the Metis system, we expressed this operon in an *E. coli* strain in which the Dam methyltransferase was deleted. As in the case of the Metis system, the MisC-MisB operon did not protect against phages when Dam was inactive, indicating that N⁶-adenine methylation of the host genome is required for defense (Figure 4C). T6 *denA* mutants that overcame defense by the MisA–MisB Metis system also displayed resistance to the MisC– MisB system, indicating that both systems require phage-mediated host genome degradation for activation (Figure S5). We therefore designate this operon type II Metis. As in the case of the MisA-MisB Metis (which we henceforth refer to as type I Metis), cells encoding type II Metis could not grow when m⁶dAMP was mixed with the growth medium. These results suggest that type II Metis is also activated by the methylated adenine nucleotide (Figure 4D).

To further investigate the molecular basis for methylated nucleotide recognition by type II Metis we used AF3 to co-fold MisC with m⁶dAMP. AF3 modeled the nucleotide with high confidence in a small pocket predicted to be formed in the intracellular portion of MisC (Figure 4E, 4F). Multiple sequence alignment of MisC homologs shows that residues predicted to form the m⁶dAMP-binding pocket are conserved (Figure S6). Point mutation in a conserved leucine residue residing in this pocket abolished type II Metis defense, confirming its importance for system activation (Figure 4G). As in the case of type I Metis, mutation in the catalytic aspartic acid of MisB did not affect the anti-phage defensive activity, showing that the MisC transmembrane protein alone can protect against phage, and suggesting that the role of MisB in this operon is to prevent MisC toxicity in the absence of phage infection (Figure 4G, S7A, S7B).

AF3 further predicted that MisC oligomerizes into an octameric circular structure that potentially forms a membrane-spanning pore (Figure 4H). Similar analysis of diverse homologs of MisC revealed that all of these are predicted to bind m⁶dAMP and form octameric, transmembrane-spanning circular structures (Figure S8). Altogether, our data suggest that MisC in type II Metis senses phage-mediated host genome degradation by binding methylated mononucleotides, and that this binding may stimulate MisC to form a pore that would impair membrane integrity.

## Discussion

Combined, our results suggest a model for Metis activity against phage propagation (Figure 5). Under normal growth conditions in the absence of phage infection, the Dam enzyme methylates GATC sequences on the double-stranded DNA polymer, so that roughly one in every 256 adenines in the genome is methylated. These methylated adenines are released to the cytosol as m^6^dAMP only in rare cases of partial endogenous DNA degradation, most likely during DNA repair, and these are rapidly “cleaned” from the cell by MisB to prevent MisA or MisC toxicity. When a lytic phage degrades the bacterial genome into individual nucleotides, the intracellular concentration of m^6^dAMP rises at least 100-fold (Figure 3D). These high concentrations of m^6^dAMP likely overwhelm the capacity of MisB to degrade all m^6^dAMP, allowing MisA activation by m^6^dAMP which leads to NAD^+^ depletion and prevention of phage propagation. Metis defense does not save the bacterium from phage-induced death, because cells cannot recover after their genome was completely degraded. Rather, Metis does not allow phages to replicate in the infected cell, thus saving neighboring bacteria from phage spread.

**Figure 5.**
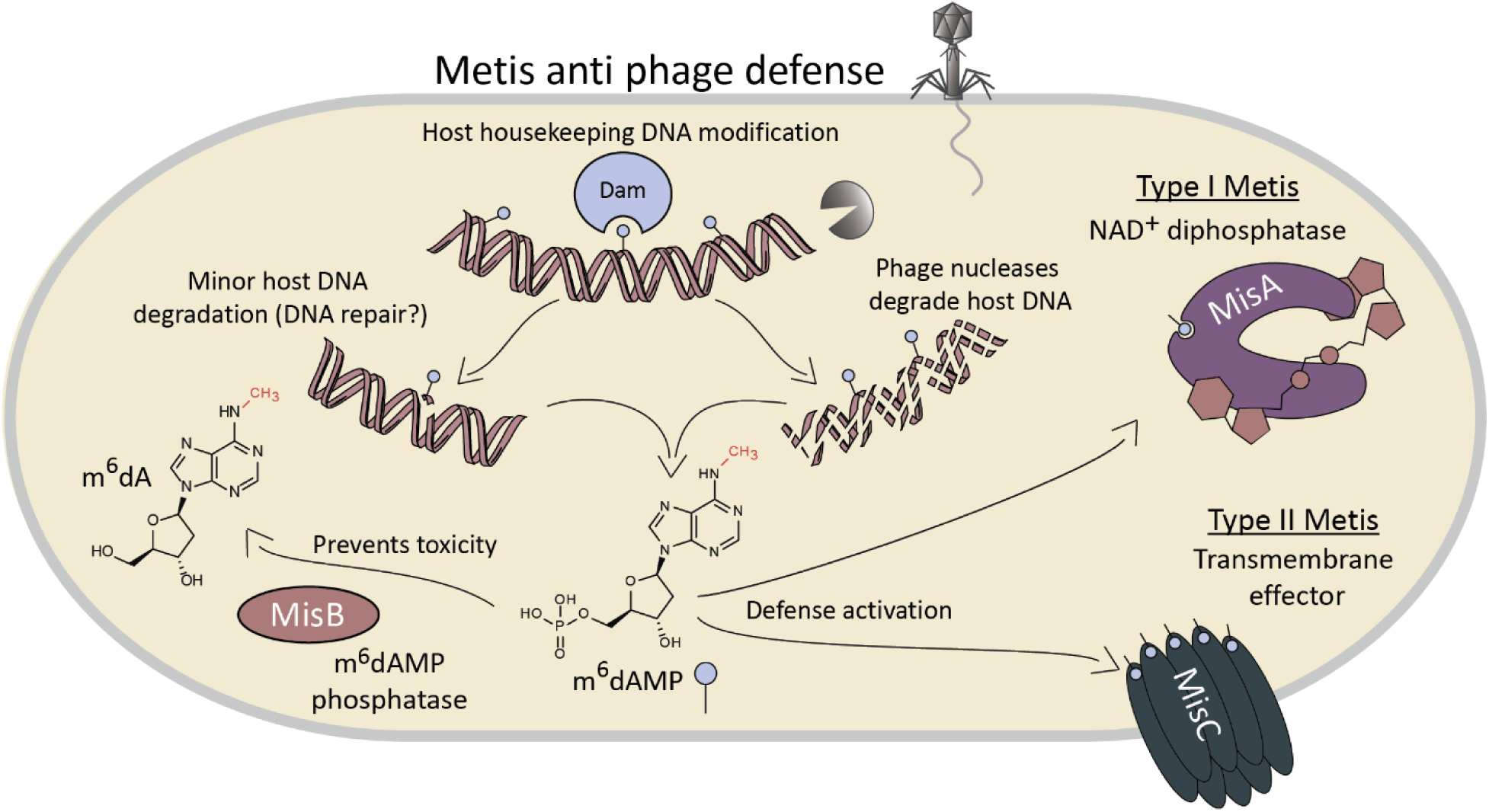
**Model for the mechanism of Metis anti-phage defense.**

Many bacterial defense systems recognize a specific phage component that triggers their defensive function^29,30^. Avs defensive proteins, for example, sense the phage portal or terminase proteins to activate defense^31,32^, and DSR2 is triggered by binding tail-tube proteins of certain phages^17,33^. While effective, this strategy limits protection to only those phages encoding the recognized proteins. By contrast, Metis provides broad defense against diverse lytic phages because it senses a consequence of phage infection - the degradation of the host genome - rather than a phage-encoded molecule. This strategy comes with its own limitation: Metis cannot defend against temperate phages, which typically do not degrade the host genome during their life cycle^34^. In addition, lytic phages can evade Metis by mutating genes required for host genome degradation (Figure 1F), although such evasion may come at a cost, as these phages lose access to host-derived nucleotides needed for efficient replication of their own genomes^35^ .

NAD^+^ depletion is a common outcome of bacterial defense systems^13^. This enzymatic activity was demonstrated in the case of Sirtuin (SIR2), TIR and SEFIR effector domains associated with many defense systems^15,17,36^. To date, all NAD^+^-depleting defensive proteins were shown to cleave NAD^+^ between the nicotinamide ring and the ribose, generating free nicotinamide and ADP-ribose (ADPR)^37^. MisA is the first anti-phage protein shown to cleave NAD^+^ in a different manner, generating the molecules NMN and AMP (Figure 2C). This form of NAD^+^ cleavage is expected to prevent phages from utilizing NAD^+^ reconstitution pathway 1 (NARP1) to alleviate defense, because NARP1 relies on ADPR and nicotinamide as substrates to rebuild NAD^+^ ^38^. The calcineurin-like metallophosphatase NAD^+^-cleaving domain found in MisA was detected in many other predicted defense systems^21^. We therefore predict that MisA-like cleavage of NAD^+^into NMN and AMP is common in bacterial defense.

While we describe here the discovery of two types of Metis systems that sense byproducts of host genome degradation, we anticipate that other types of Metis exist in nature. Recognition of methylated cytosines, for example, could also be efficient for genome degradation sensing, as cytosine methylation is a widespread epigenetic DNA modification in bacteria^39^. Furthermore, we envision that some Metis systems would encode their own DNA-modifying enzyme, so that their activity will not depend on endogenous Dam enzymes. In these systems, the DNA-modifying enzyme would install a specific base modification on the bacterial DNA, and the Metis effector protein would be triggered when this modified base accumulates as a consequence of phage-induced host genome degradation. Indeed, we detected multiple homologs of MisA that are found in an operon with DNA adenine methyltransferases or other predicted methylases, forming putative type III Metis systems (Figure S9, Supplementary Table S3).

Many principles of antiviral immunity are shared between bacterial and eukaryotic immune pathways, and some components of the human immune arsenal have been evolutionarily derived from bacterial and archaeal defense systems^14,40,41^. Viruses that infect eukaryotes do not commonly degrade the nuclear genomes, although some viruses, including those infecting freshwater algae, shred the genome of the infected cell as part of their life cycle^42^, and some animal viruses, for example frog-infecting Ranaviruses, are also thought to cause host DNA degradation^43^. As cytosines are frequently methylated in eukaryotic genomes in the context of CpG motifs^44^, it might be possible that defense pathways in eukaryotes can manifest similar principles as Metis, and monitor the integrity of the cellular genome by sensing methylated single-nucleotide cytosines. Whether such immune pathways indeed exist in eukaryotes remains to be determined.

## Acknowledgements

We thank members of the Sorek lab for constructive discussions during this study. R.S. was supported, in part, by the European Research Council (grant ERC-AdG GA 101018520), the Israel Science Foundation (MAPATS grant 2720/22), the Deutsche Forschungsgemeinschaft (SPP 2330, grant 464312965), the Minerva Foundation with funding from the Federal German Ministry for Education and Research, a research grant from the Estate of Hermine Miller, the Center for Immunotherapy at the Weizmann Institute of Science, and the Knell Family Center for Microbiology. I.O. was supported by the Ministry of Absorption New Immigrant program. E.Y. was supported by the Clore Scholars Program and, in part, by the Israeli Council for Higher Education (CHE) via the Weizmann Data Science Research Center.

## Materials and Methods

### Strains and growth conditions

All *E. coli* strains were grown in MMB media (lysogeny broth (LB) supplemented with 0.1 mM MnCl2 and 5 mM MgCl2) at 37°C with 200 rpm shaking or on solid 1.5% LB agar plates. Ampicillin 100 μg/mL, chloramphenicol 30 μg/mL or kanamycin 50 μg/mL were added when necessary for plasmid maintenance. *E. coli* DH5a (NEB) was used for cloning, BL21 (DE3) for protein purification and MG1655, BW25113 JW3350 (Δ*dam*), and JW1944 (Δ*dcm*) for experiments with phages. All chemicals were obtained from Sigma Aldrich unless stated otherwise. All phages used in the study were amplified from a single plaque (at 37°C in *E. coli* MG1655 culture in MMB until the culture collapsed. A list of all plasmids, strains and phages used in this study can be found in Supplementary Table S3.

### Plasmid construction and transformation

DNA amplification for cloning was performed using KAPA HiFi HotStart ReadyMix (Roche) according to the manufacturer’s instruction. All primers were obtained from Sigma Aldrich. Supplementary Table S2 lists all primers used in this study.

For cloning of large fragments, PCR products with 20-nucleotide overlaps were generated and treated with FastDigest DpnI restriction enzyme (ThermoFisher) for 30 min at 37°C. Fragments were then Gibson-assembled by NEBuilder HiFi DNA Assembly Master Mix (NEB) according to the manufacturer’s instructions and used for transformation into DH5a (NEB #C2987H). Single colonies were checked by PCR and plasmids were validated by a plasmid sequencing service (Plasmidsaurus). Plasmids ordered from Twist Bioscience or GenScript Corporation are listed in Supplementary Table S3. Verified plasmids were used for the transformation of *E. coli* strains using the standard TSS protocol.^45^

### Plaque assays

Phage titer was determined as described previously^46^. 300 µL of the overnight bacterial cultures were mixed with 30 mL of melted MMB 0.5% agar, poured on 10 cm square plates and left to dry for 1 h at room temperature. L-Arabinose was added to a concentration of 0.2% to induce gene expression from pBAD plasmids. IPTG was added to a concentration of 1 mM to induce expression from pBba6c plasmid. Tenfold dilutions of phages were prepared in MMB and 10 µL of each dilution was dropped onto the plates. Plates were incubated overnight at 25°C (for type II Metis) and 37°C (for type I Metis). Plaque-forming units were counted the next day.

### T4 and T6 escapers selection and DNA sequencing

10^8^-10^9^ PFU of T4 and T6 were spread on Petri dishes on which cells expressing Metis 401675 were grown. On the next day, individual plaques were collected, and isolated using three transfers on *E. coli* MG1655 cells. Isolated phages were tested via plaque assays on cells expressing Metis 401675 and resistant phages were selected for DNA purification and genome sequencing.

DNA from escaper phages was extracted from 500 µL of a high-titer phage lysate (> 10^7^ PFU/mL) as described previously^29^. Phage genomes sequencing was performed by Plasmidsaurus. Wild-type (WT) T4 and T6 genomes were assembled with SPAdes v3.15.3 (default parameters). Escaper reads were analyzed with breseq v0.38.1 (default parameters) using the corresponding WT phage genome as reference. Data on isolated escapers are found in Supplementary Table S4.

### Liquid infection assay

Overnight bacterial cultures of *E. coli* MG1655 cells expressing Metis from *E. coli* 401675 and control *E. coli* MG1655 cells with an empty vector were diluted in MMB (1:100) and grown until reaching an optical density at 600nm (OD_600_) of 0.3 at 37°C. Then, 180 µL of cultures were transferred to a 96-well plate and infected with 20 µL of phages at various MOIs. Culture growth was followed by OD_600_ measurements every 10 min on a Tecan Infinite 200 plate reader at 37°C.

### Liquid growth and toxicity assay

Overnight bacterial cultures were diluted in MMB (1:100), supplemented (or not) with m^6^dAMP (50 µM for type I Metis and 500 µM for type II Metis) and grown in 20 µL in a 384-well plate at 37°C. following by OD_600_ measurements every 10 min on a Tecan Infinite 200 plate reader.

### Cell lysates preparation

*E. coli* MG1655 cells carrying a plasmid with the Metis 401675 system, MisA alone, MisB alone or an empty plasmid were grown at 37 °C, 200 rpm until reaching an OD_600_ of 0.3 in 150 mL of MMB. Cells were then infected with T7 phage with MOI=3. Samples were collected before infection and 15 min after infection. At each time point, 50 mL of cells were centrifuged for 10 min at 4°C, 4000 *g* for 10 min, and the pellet was frozen in liquid nitrogen and stored at −80°C. To extract cell metabolites from frozen pellets, 600 μL of 100 mM sodium phosphate buffer (pH 7.5) was added to each pellet. Samples were transferred to FastPrep Lysing Matrix B in a 2 mL tube (MP Biomedicals, cat #116911100) and lysed at 4°C using a FastPrep bead beater for two rounds of 40 s at 6 ms^−1^. The tubes were then centrifuged at 4°C for 10 min at 15,000 *g*. The supernatant was then transferred to an Amicon Ultra-0.5 Centrifugal Filter Unit 3 kDa (Merck Millipore, no. UFC500396) and centrifuged for 45 min at 4°C, 12,000 *g*. Lysates were analyzed by LC-MS.

### LC-MS analysis of the lysates

Metabolic profiling of polar metabolites within filtered cell lysates was carried out as described previously^47^ with minor modifications as described below. In brief, analysis was carried out using an Acquity I class UPLC System combined with a mass spectrometer Q Exactive Plus Orbitrap (Thermo Fisher Scientific), which was operated in positive ionization mode. The LC separation was carried out using the SeQuant Zic-pHilic (150 mm × 2.1 mm) with the SeQuant guard column (20 mm × 2.1 mm; Merck). Mobile phase B was acetonitrile and mobile phase A was 20 mM ammonium carbonate with 0.1% ammonia hydroxide in deionized distilled water/acetonitrile (80:20, v/v). The flow rate was kept at 200 μL/min, and the gradient was as follows: 0–2 min 75% of B, 14 min 25% of B, 18 min 25% of B, 19 min 75% of B, for 4 min, 23 min 75% of B. Peak areas were extracted using MZmine 2 with an accepted deviation of 5 ppm. All molecules used as standards were obtained from Sigma.

### NAD^+^ detection assay

*E. coli* MG1655 cells carrying a plasmid with the Metis 401675 system or an empty plasmid were grown at 37 °C, 200 rpm until reaching an OD_600_ of 0.3, and then supplemented with 50 µM of m⁶dAMP. Plates were incubated at 37°C with shaking in a TECAN Infinite200 plate reader. 15 µL of cells were taken before molecule supplementation (t=0) and at 15, 30 and 60 min post-supplementation. Cells were mixed with 20 µL of 100% ethanol and kept at −20°C. For the NAD^+^ detection assay, samples were diluted 1:5 in 100 mM sodium phosphate buffer (pH 7.5). Then 5 µL of the samples were mixed with 5 µL of NAD/NADH-Glo™ detection reagent (Promega corp.) in a Nunc^®^ 384 well polystyrene plate (Sigma p5991-1CS). Plates were incubated at 25°C in a TECAN Infinite200 plate reader and luminescence was measured every 10 min.

### Protein purification

MisA (Metis 402837) and MisB (Metis 401675) were cloned into the pET28-His-bdSUMO expression vector (TWIST Bioscience, USA), which encodes an N-terminal His14-bdSUMO tag^48^. Plasmids were transformed into LOBSTR-BL21(DE3)-RIL cells (Kerafast, USA), and cells were grown in 50 mL of MMB media at 37°C until mid-log phase (OD_600_∼0.5), then IPTG was added to a concentration of 0.5 mM and cells were further grown overnight at 16°C, centrifuged and flash-frozen. His-tagged proteins were purified using 50 µL NEBExpress® Ni-NTA Magnetic Beads (NEB) according to the manufacturer’s protocol. Elution was done using 1 µg His-tagged bdSENP1 protease in 100 µL of cleavage buffer (20 mM HEPES 125 KCl and 1 mM DTT), for 12 hours at 4°C with intensive shacking.

### Generation of m⁶dAMP and m⁶AMP

m⁶dATP and m⁶ATP from Jena Bioscience was diluted in Apyrase buffer (NEB) to 10mM and incubated with 1 unit of Apyrase (NEB) in 50 µL for 1 hour at 30°C. The efficiency of cleavage was analyzed by HPLC and LC-MS.

### Enzymatic activity of MisA

100 µM of NAD^+^ or ATP were incubated with 1 µM of the purified MisA in 10 mM Tris-HCl buffer (pH=7) with supplementation of 10 µM synthetic adenine variants for 120 min at 37°C. The products of the reactions were diluted 1:200 in 100 mM sodium phosphate buffer (pH 7.5) and were mixed with 5 µL of NAD/NADH-Glo™ detection reagent (Promega corp.) or BacTiter-Glo™ Microbial Cell Viability Assay in a Nunc^®^ 384 well polystyrene plate (Sigma p5991-1CS). Plates were incubated at 25°C in a TECAN Infinite200 plate reader and luminescence was measured every 10 min. Priore to HPLC analysis the products of the reactions were diluted 1:4 in water and were transferred to an Amicon Ultra-0.5 Centrifugal Filter Unit 3 kDa (Merck Millipore, no. UFC500396) and centrifuged for 45 min at 4 °C, 12,000 *g.* dAMP was obtained from Sigma, m⁶dATP from Jena Bioscience, m⁶dA from Angene, m⁶adenine from Cayman, and m⁶dAMP and m⁶AMP were generated using Apyrase treatment as described above.

### HPLC analysis of the MisA *in vitro* reactions

20 µL of the reactions obtained in the previous step after filtration were analyzed using HPLC. HPLC of the obtained fraction was performed using Agilent 1260 and chromatography SUPELCOSIL™ LC-18-T HPLC Column. The following protocol was used for all runs: 1 min of mobile phase A 100%, 2 min 75% A and 25% B, 2 min 50% A and 50% B, 2 min 20% A and 80% B and 3 min 100 % A, 1 mL/min flow rate. Mobile phase A was 20 mM potassium phosphate pH 6 and B was 20 mM potassium phosphate pH 6 in 20% methanol.

### Enzymatic activity of MisB

100 µM of m⁶AMP was incubated with 1 µM of the purified MisB in 10 mM Tris-HCl buffer (pH=8) with supplementation of 10 mM of MgCl_2_ for 120 min at 37°C. The products of the reactions were diluted 1:4 in water and were transferred to an Amicon Ultra-0.5 Centrifugal Filter Unit 3 kDa (Merck Millipore, no. UFC500396), centrifuged for 45 min at 4°C, 12,000 *g*, and analyzed by LC-MS on Waters SYNAPT-XS Q-Tof mass spectrometer.

### LC-MS analysis of the *in vitro* reactions

The sample solutions were analyzed by UPLC–HRMS without dilution. The analyses were carried out on a Waters SYNAPT-XS Q-Tof mass spectrometer (Manchester, UK) with an electrospray ionization (ESI) source.

The spectra m^6^dAMP and m^6^dA were recorded in the negative ion mode within a mass range from 100 to 1200 m/z. The parameters were set as follows: capillary voltage at 1.5 kV, cone gas flow at 50 L/h, source temperature at 140°C, and cone voltage at 20 V (−). The desolvation temperature was set at 600°C, and the desolvation gas (N2) flow rate was set at 800 L/h.

Lock spray was acquired with Leucine Encephalin (m/z=554.2615 in negative mode) at a concentration of 200 ng/mL and a flow rate of 10 μL/min once every 10 s for 1 s period to ensure mass accuracy. Waters MassLynx v4.2 software was used for data acquisition and data processing.

The analytes were separated using Waters Premier Acquity UPLC system. The gradient elution was achieved with Waters Acquity Premier HSS T3 Column, 1.8 μm, 2.1 × 100 mm at 0.4 mL/min flow rate, 35°C. Mobile phase A consisted of 20 mM aqueous Ammonium Acetate (Sigma-Aldrich 09691) and mobile phase B consisted of 20 mM Ammonium Acetate in acetonitrile: water 75:25. The gradient was as follows: 0–2.5 min 0% of B, 2.5-6 min 10% of B.

### Structure prediction

AlphaFold3^24^ with default parameters was used to generate structural models for MisA, MisB and MisC together with the respective ligands. The following SMILES code were used for the ligands: with default parameters was used to generate structural models for MisA, MisB and MisC together with the respective ligands. The following SMILES code were used for the ligands:

NAD^+^:O[C@H]([C@@H]1O)[C@@H](O[C@@H]1COP(OP(OC[C@H]([C@@H](O)[C@H]2O)O[C@H]2[N+]3 =CC(C(O)=O)=CC=C3)(O)=O)(O)=O)N(C=N4)C5=C4C(N)=NC=N5

m⁶dAMP: O[C@H]1C[C@@H](O[C@@H]1COP(O)(O)=O)N(C=N2)C3=C2C(NC)=NC=N3

### Phylogenetic analysis of MisB homologs

Protein sequences of all genes in 38,167 bacterial and archaeal genomes were downloaded from the IMG database in October 2017 and clustered into groups of homologs as previously described^36^. Protein sequences of genes from metagenome scaffolds were downloaded from the IMG database in April 2020 and merged into the set of protein clusters from isolate genomes as previously described^41^.

All non-redundant homologs of MisB were collected from its cluster and listed in the Supplementary table S1. The protein sequences were aligned using MAFFT (v7.520)^49^ with default parameters and mode auto. The phylogenetic tree was constructed using IQ-TREE (v2.2.0)^50^. Selection of the best-fit model of amino acid substitution was inferred using the ModelFinder function in IQ-TREE^51^, with model Q.pfam+I+G4 chosen according to Bayesian Information Criterion. Node support was computed using 1,000 iterations of the ultrafast bootstrap function in IQ-TREE (option -bb 1000)^52^. All trees were visualized with iTOL^53^.

Leaves of the phylogenetic tree were annotated based on the presence of MisA or MisC, if a MisA or a MisC homolog was detected within two genes upstream or downstream of MisB. MisA was identified by the presence of calcineurin-like phosphoesterase domain (Pfam PF00149), and MisC was identified by the presence of transmembrane regions. Transmembrane regions and the number of transmembrane helices were predicted using TMHMM^54^.

## Supplementary Figures

**Supplementary Figure S1.**
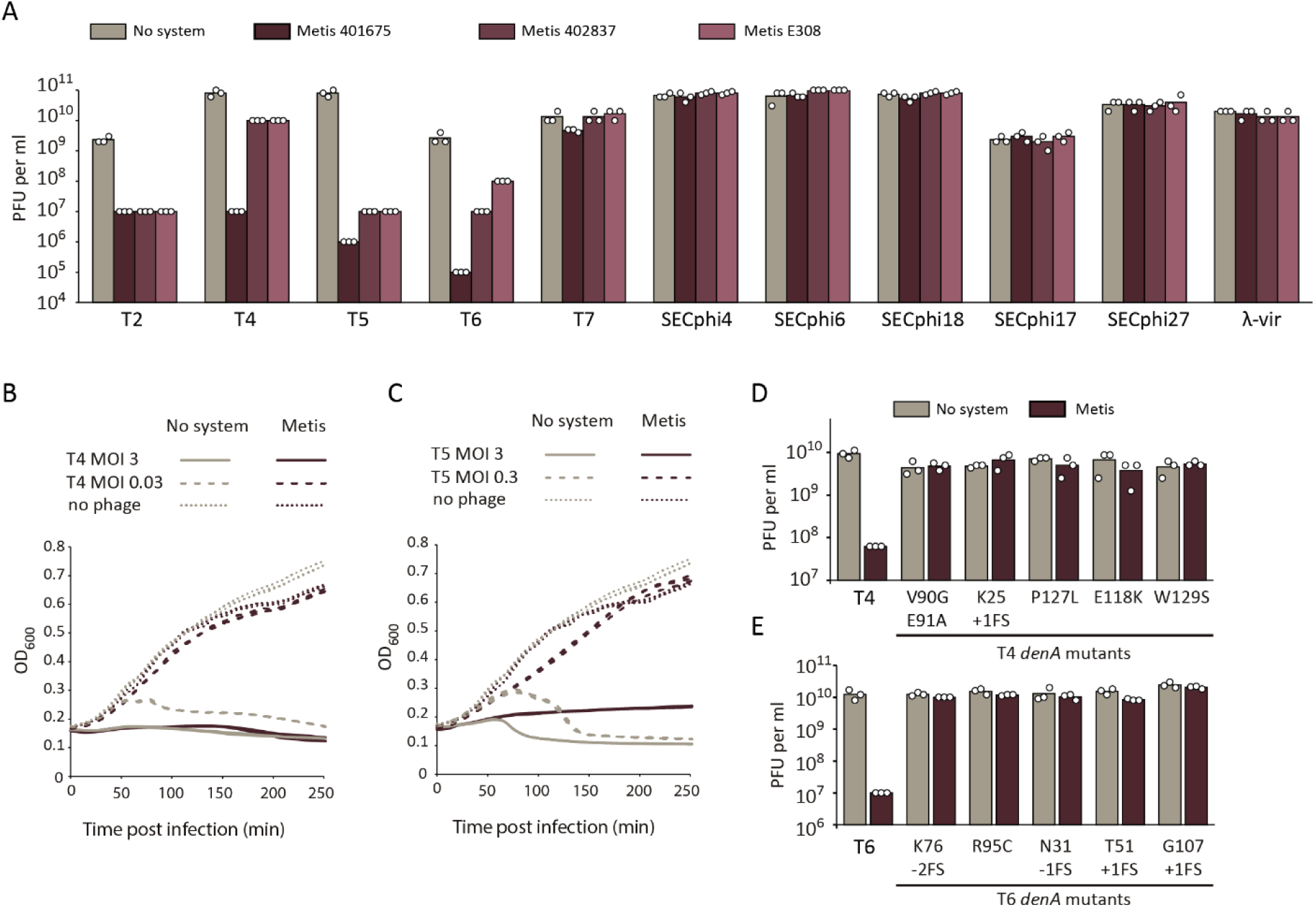
A. Serial dilution plaque assay experiments. Data represent PFUs per milliliter of phages infecting control cells (no system) and cells expressing Metis systems cloned from *E. coli* strains 401675, 402837, and E308 and expressed in *E. coli* MG1655. Bar graphs represent the average of three independent replicates, with individual data points overlaid. B, C. Growth curves of *E. coli* MG1655 cells expressing the Metis system from *E. coli* 401675, or an empty vector as control, infected by phage T4 (B) at an MOI of 0.03 or 3 or phage T5 (C) at an MOI of 0.3 or 3. Data from three replicates are presented as individual curves. D. T4 escaper mutants. E. T6 escaper mutants. Panels D and E both show plating efficiency of WT or mutated phages on *E. coli* MG1655 cells (light gray), or on *E. coli* MG1655 cells expressing the Metis system (dark red). Data represent plaque-forming units (PFUs) per milliliter. Bar graphs represent the average of three independent replicates, with individual data points overlaid.

**Supplementary Figure S2.**
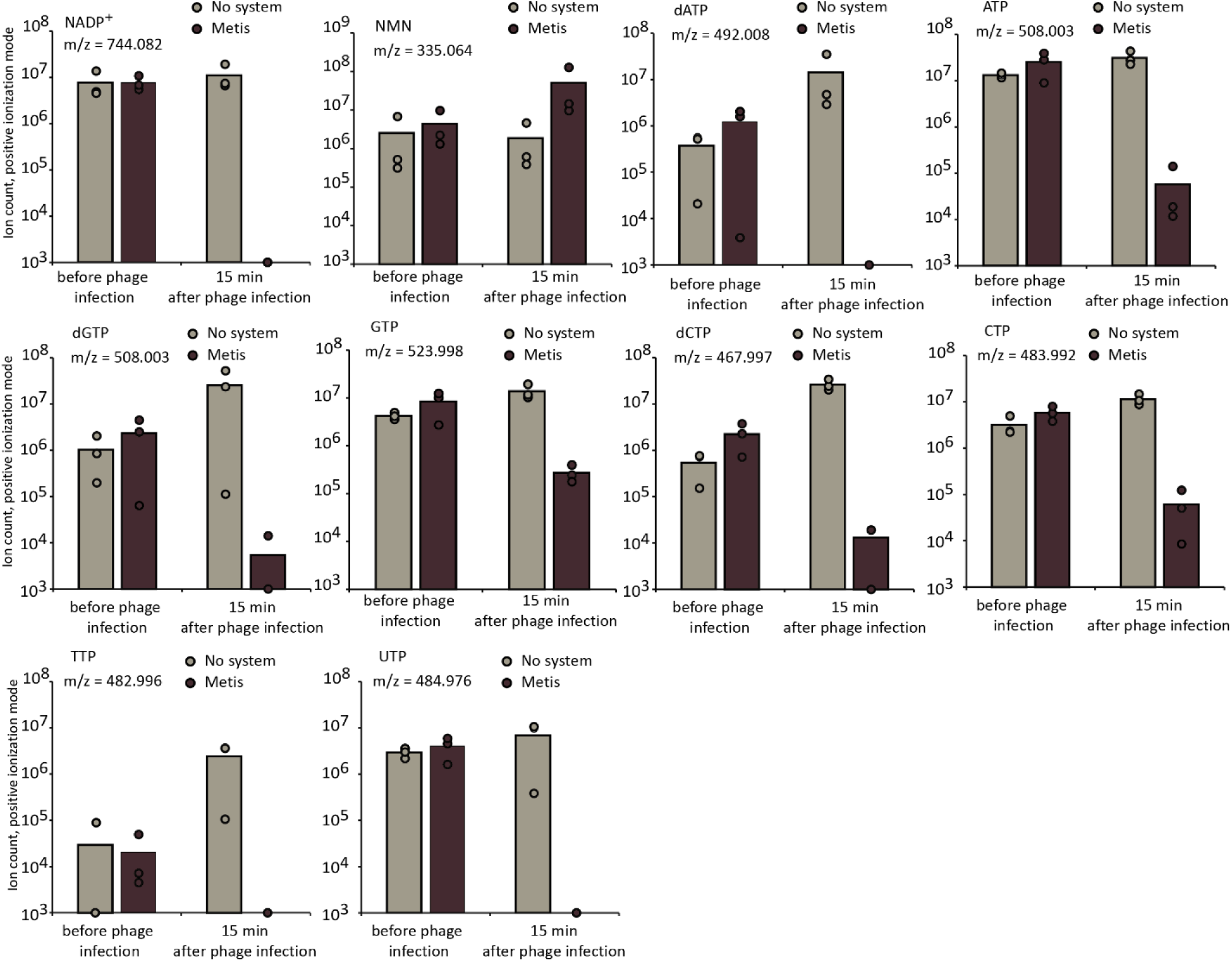
Metabolites affected by Metis activity. Shown are ion counts for selected metabolites in uninfected cells or in cells 15 minutes after infection with T7 phage (MOI = 3). Presented are liquid chromatography LC-MS ion count data; bars represent the mean area under the curve of three experiments, with individual data points overlaid. Values of m/z are indicated within the panels; identification of the molecules was made by comparing with pure standards.

**Supplementary Figure S3.**
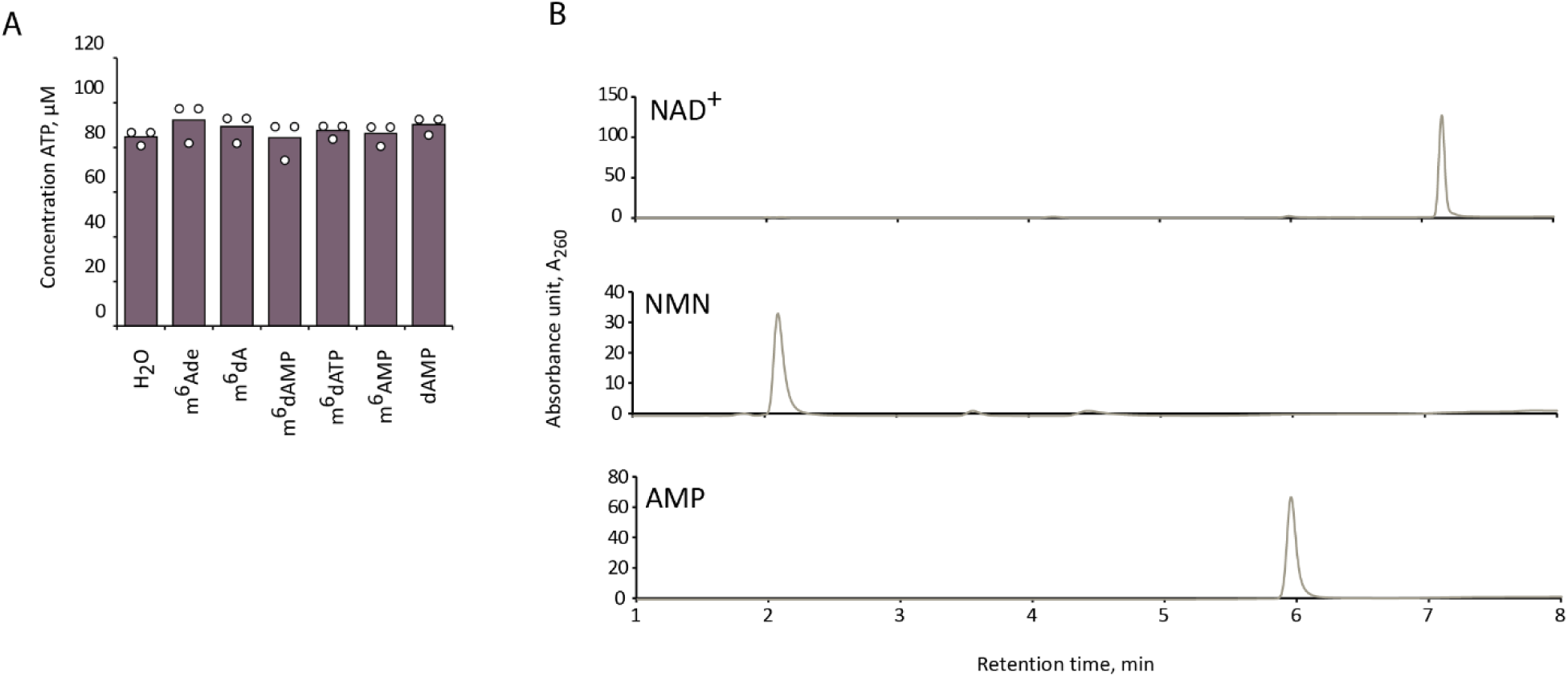
A. MisA does not degrade ATP *in vitro* in the presence of m^6^dAMP. 100µM ATP was incubated with 1 µM purified MisA and 10 µM synthetic adenine variants for 120 minutes, and ATP levels were measured using the BacTiter-Glo biochemical assay (Methods). Average of three replicates with individual data point overlaid. B. HPLC analysis of chemical standards of NAD^+^, NMN and AMP.

**Supplementary Figure S4.**
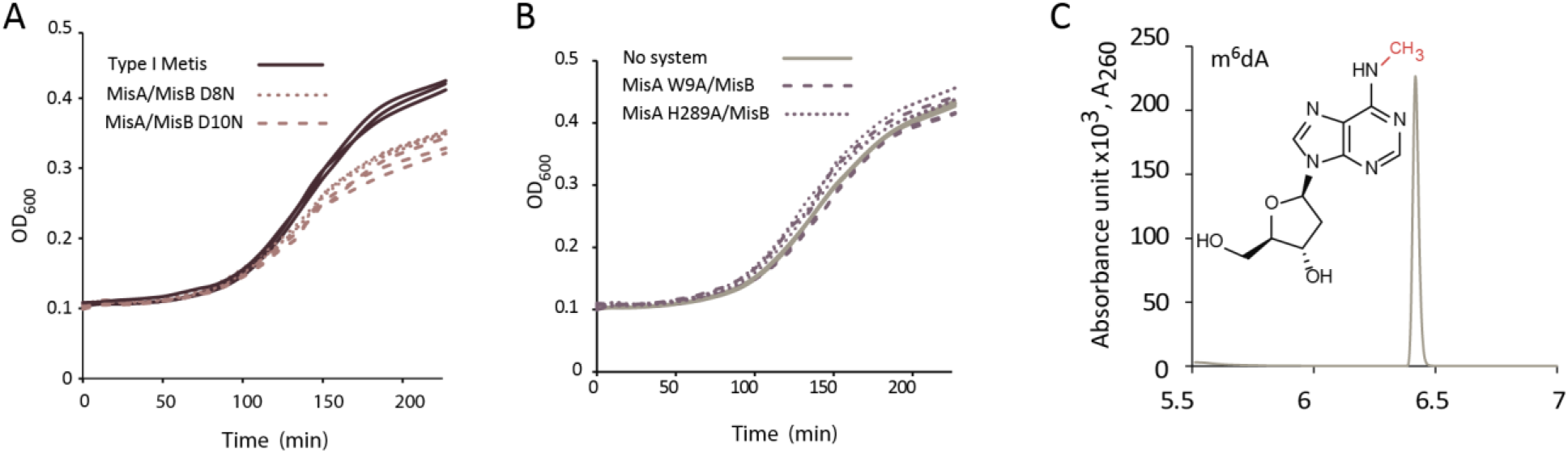
A. Growth curves of *E. coli* MG1655 cells expressing WT Metis or Metis with point mutations in MisB. B. Growth curves of *E. coli* MG1655 cells expressing WT Metis, Metis with mutated MisA, or empty vector. Data from three replicates are presented as individual curves. C. LC-MS analysis of m^6^dA chemical standard.

**Supplementary Figure S5.**
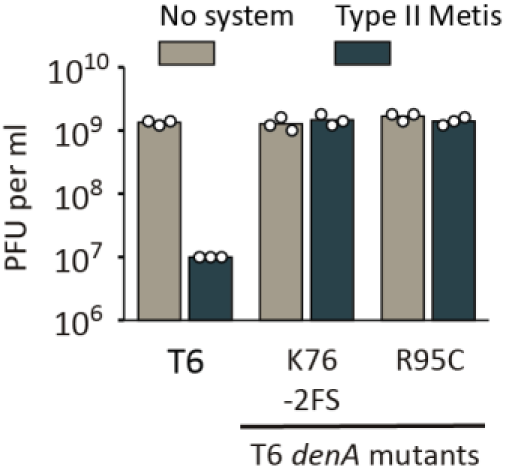
T6 *denA* mutants, which escape type I Metis, also escape type II Metis defense. Shown is plating efficiency of WT or mutated phages on *E. coli* BW25113 cells (light gray), or on *E. coli* BW25113 cells expressing the type II Metis system (dark green). Data represent plaque-forming units (PFUs) per milliliter. Bar graphs represent the average of three independent replicates, with individual data points overlaid.

**Supplementary Figure S6.**
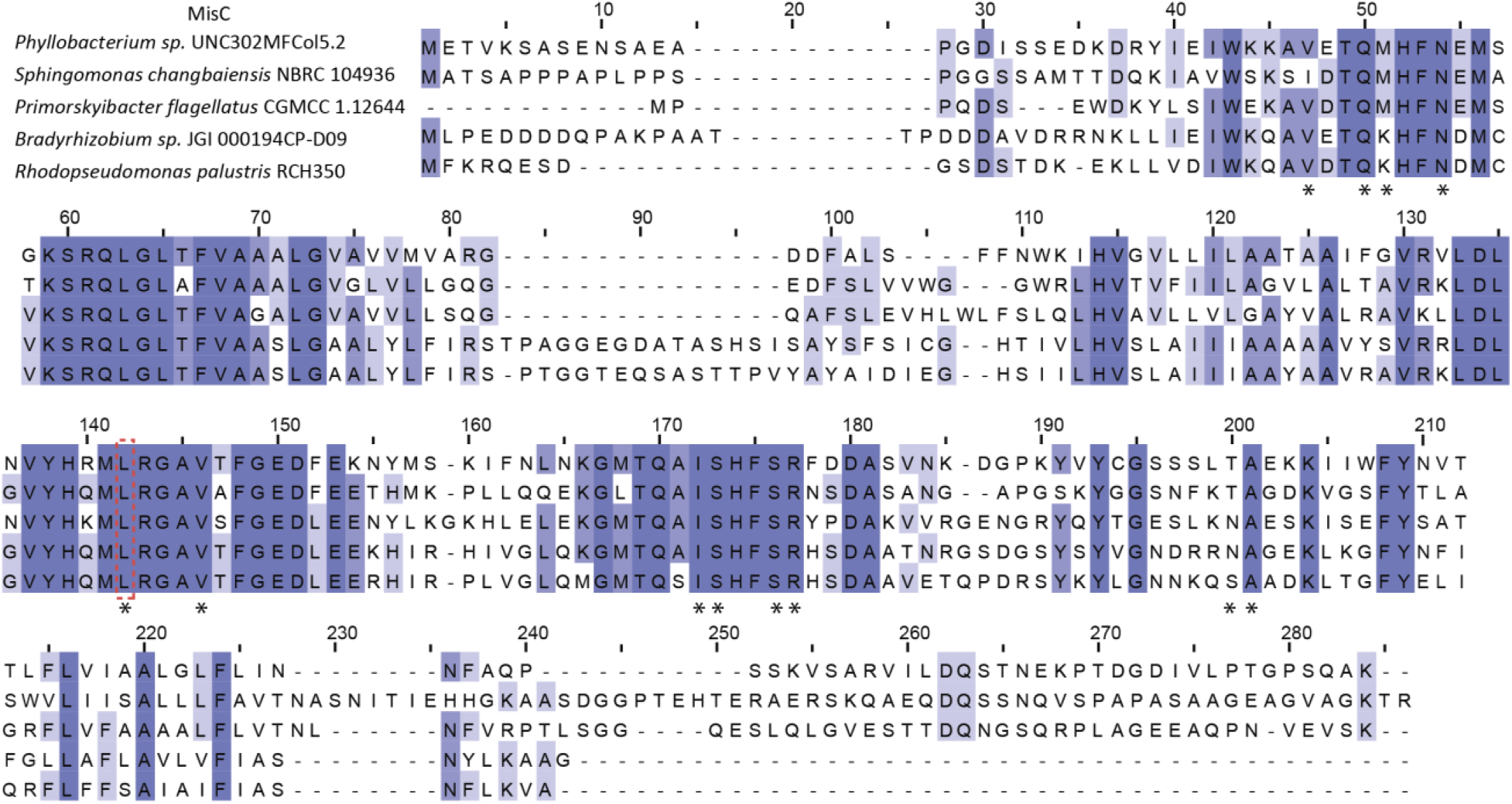
Sequence alignment of MisC proteins. Conserved residues are in purple. Residues present in the m^6^dAMP binding pocket (defined as residues with a distance of up to 4 Å from m^6^dAMP in the predicted AF3 structure) are marked with asterisks. Red square indicates residue mutated in this study.

**Supplementary Figure S7.**
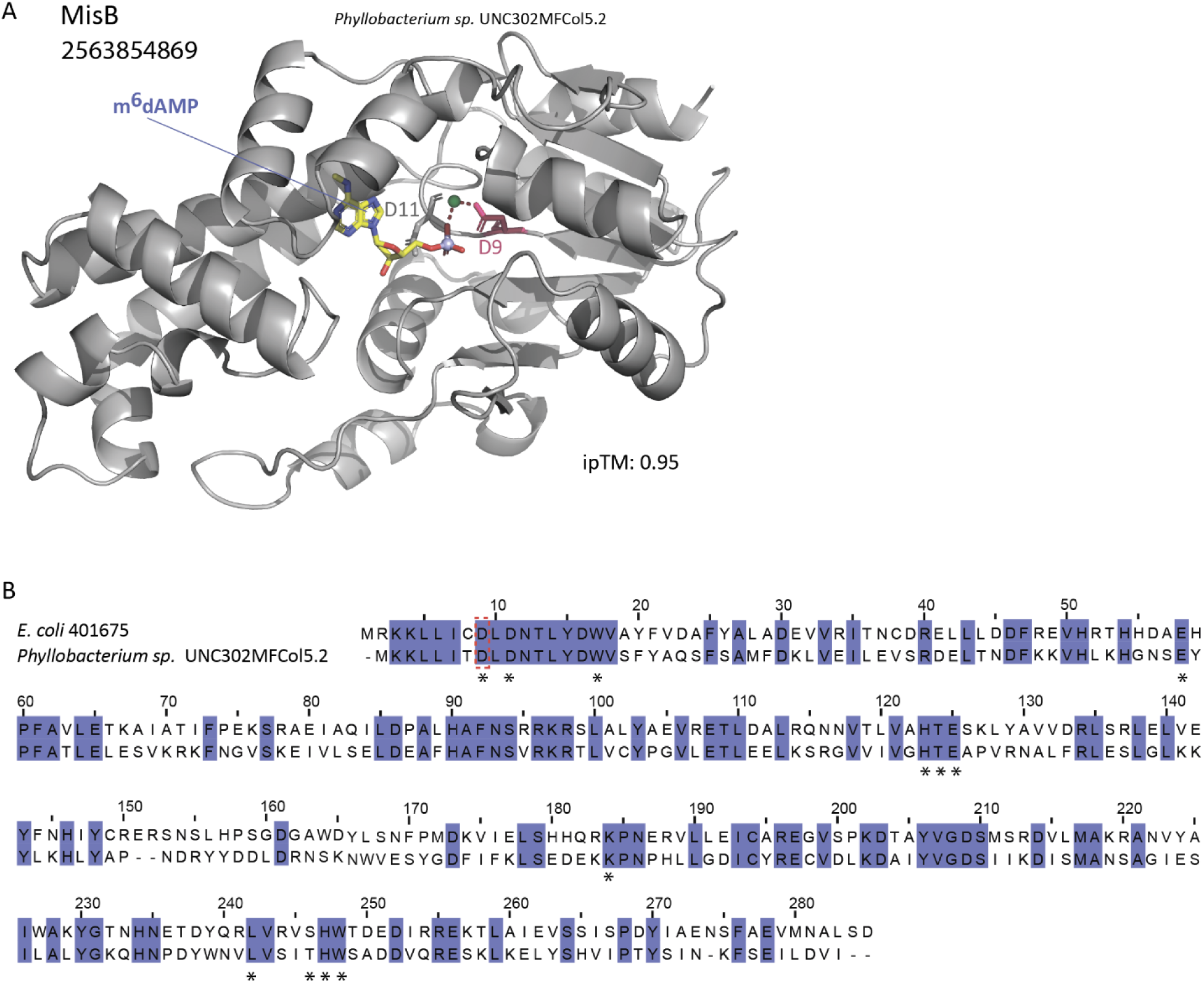
A. AlphaFold3-predicted structures of MisB from *Phyllobacterium sp.* UNC302MFCol5.2 in complex with m⁶dAMP and Mg^2+^ (iPTM = 0.95). Residue mutated in this study is presented in pink, Mg^2+^ ion is green. Red dashes indicate predicted hydrogen bonding interactions. B. Sequence alignment of MisB proteins from *E. coli* 401675 (type I Metis) and *Phyllobacterium sp.* UNC302MFCol5.2 (type II Metis). Conserved residues are in purple. Residues present in the m^6^dAMP binding pocket (defined as residues with a distance of up to 4 Å from m^6^dAMP in the predicted AF3 structure) are marked with asterisks. Red square indicates residue that was mutated in this study.

**Supplementary Figure S8.**
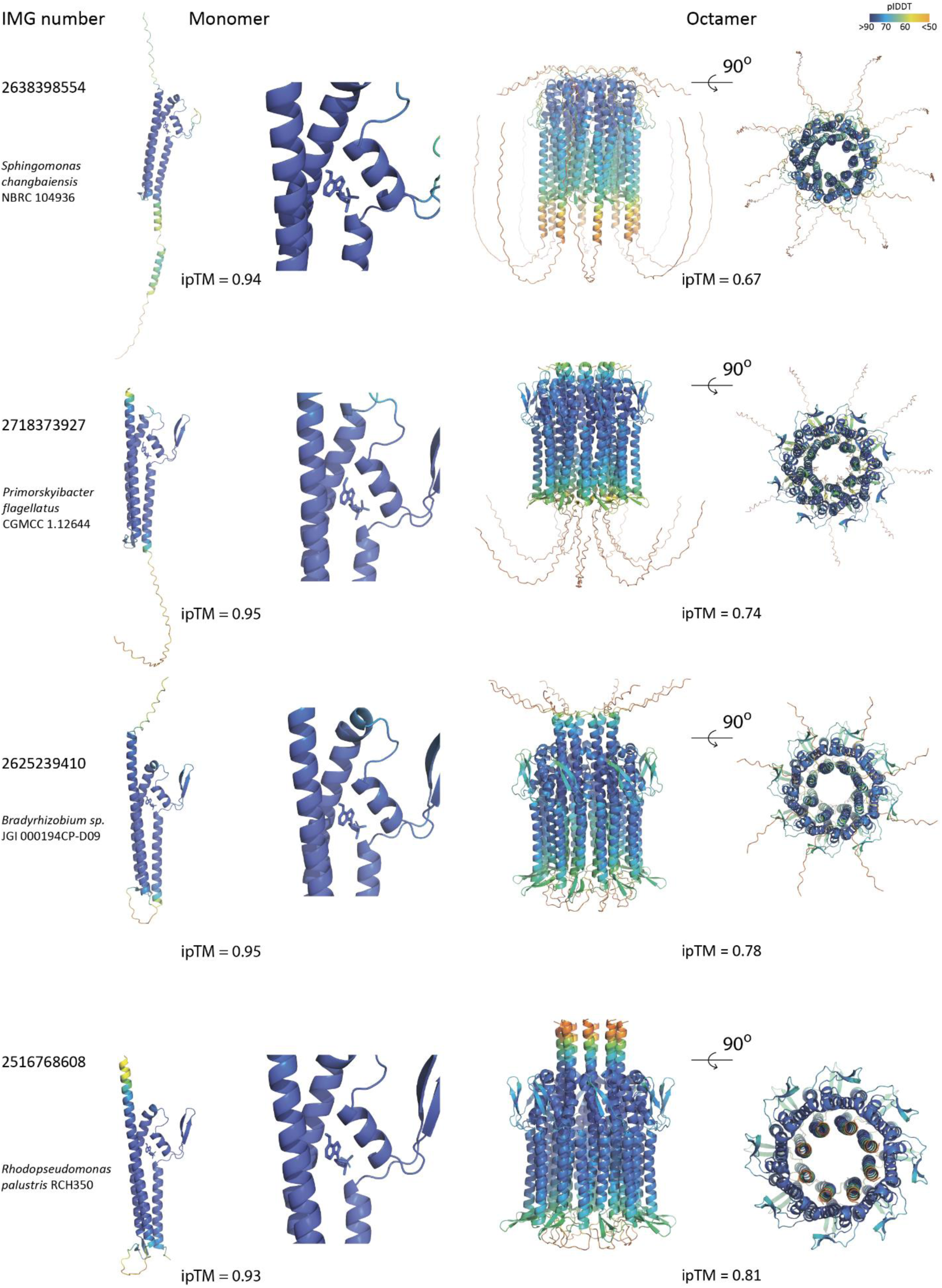
AlphaFold3-predicted structures of MisC homologs in monomeric forms co-folded together with m^6^dAMP, and the corresponding predicted octameric forms. Colors represent pLDDT scores. Numbers on the left represent the gene accession in the IMG database.

**Supplementary Figure S9.**
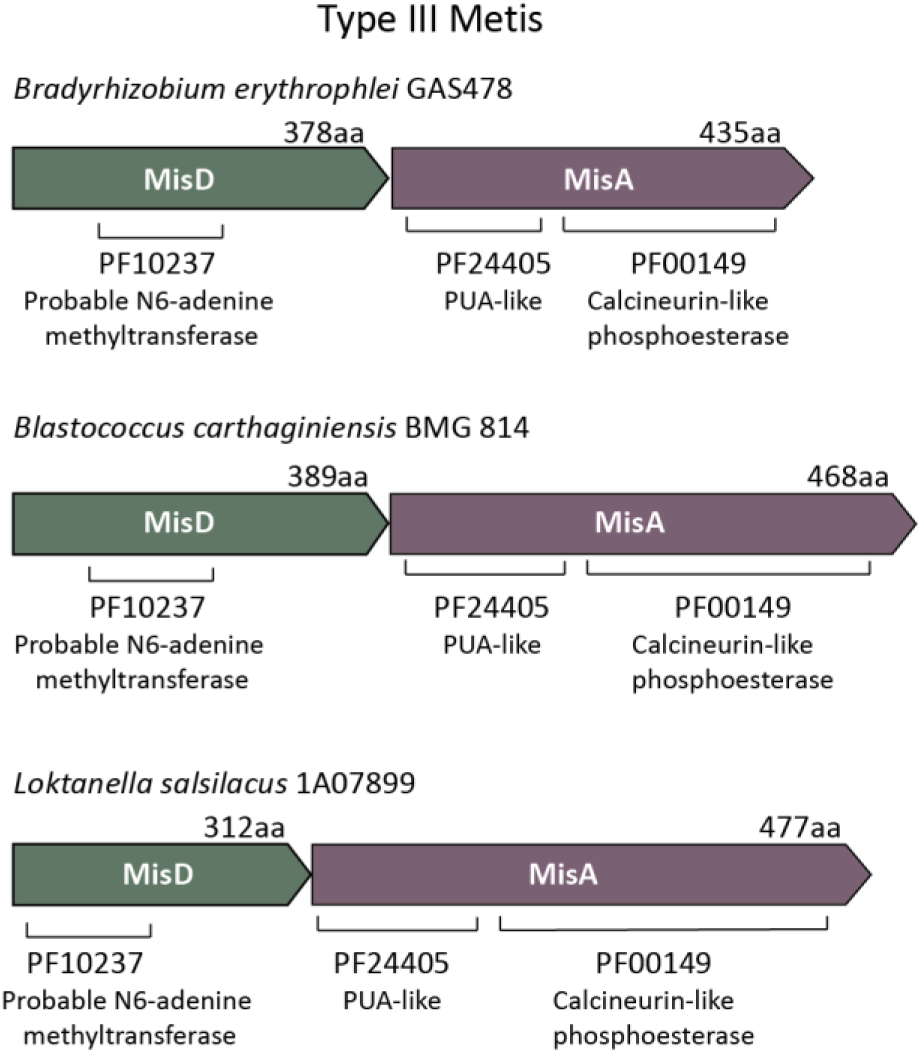
Predicted type III Metis systems. Shown are operons in which a MisA homolog is associated with a gene annotated as encoding a methyltransferase. These operons are hypothesized to comprise an anti-phage defense similar to type I Metis, but in which the associated methylase (called here MisD) methylates the host genome rather than the host-encoded Dam enzyme.

